# Weak epistasis may drive adaptation in recombining bacteria

**DOI:** 10.1101/119958

**Authors:** BJ Arnold, MU Gutmann, YH Grad, SK Sheppard, J Corander, M Lipsitch, WP Hanage

## Abstract

The impact of epistasis on the evolution of multilocus traits depends on recombination. Population genetic theory has been largely developed for eukaryotes, many of which recombine so frequently that epistasis between polymorphisms has not been considered to play a large role in adaptation and has been compared to the fleeting influence of non-heritable effects. Many bacteria also recombine, some to the degree that their populations are described as ‘panmictic’ or ‘freely recombining’. However, whether this recombination is sufficient to limit the ability of selection to act on epistatic contributions to fitness is unknown. We create a sensitive method to quantify homologous recombination in five bacterial pathogens and use these parameter estimates in a multilocus model of bacterial evolution with additive and epistatic effects. We find that even for highly recombining species (e.g. *Streptococcus pneumoniae* or *Helicobacter pylori*), selection may act on the cumulative effects of weak (as well as strong) interactions between distant mutations since homologous recombination typically transfers only short segments. Furthermore, whether selection acts more efficiently on physically proximal loci depends on the average recombination tract length. Epistasis may thus play an important role in the adaptive evolution of bacteria and, unlike in eukaryotes, does not need to be strong, involve near loci, or require specific metapopulation dynamics.

## Introduction

Epistasis for fitness traits may arise from any nonlinearity in the multi-locus genotype-to-fitness map (*1*,*2*), which includes, but is not limited to, traits controlled by protein-protein interactions. The role of epistasis in adaptive evolution has been debated since the origin of population genetics (*3–7*). Fisher and other geneticists have argued that selection primarily acts on the additive effects of individual alleles (*3*,*7*,*8*), since crossover recombination in eukaryotes rapidly destroys allele combinations, thus opposing selection on epistatic effects between alleles. Exceptions to their view arise only when epistatic effects are exceedingly strong or interacting loci are tightly linked, preventing recombination (*9*). Alternatively, Wright and others have argued that species consist of metapopulations with dynamics that limit the ability of recombination to dominate microevolutionary processes, since only individuals within the same subpopulation may exchange alleles (*5*). Since evidence for these dynamics in nature is limited (*7*,*10*), epistasis has not been given a prominent role in adaptation theory.

However, these debates have focused on sexual eukaryotes, and the majority of life on earth does not sexually reproduce. Bacteria, which have colonized almost every conceivable ecological niche, also recombine to varying degrees through multiple mechanisms, leading to a continuum of genealogical structures from clonal to “fully sexual” (*11*). For microbes that recombine in moderation (e.g. *S. aureus*), it is clear that selection can easily act on epistatic allele combinations: mutations will likely exist only in the genetic background on which they arose, such that any background-specific epistatic effects are heritable through time and may spread through populations via selection. However, we do not know whether highly recombining bacteria, such as those that have little genetic linkage and have been historically labeled as “fully sexual” or “freely recombining” (*11*,*12*), decouple interacting mutations frequently enough to prevent selection on *weak* epistatic effects, and how strong epistasis must be to dominate the microevolutionary process and drive adaptation in these species. It also remains unknown if, as in eukaryotes, selection acts more efficiently on epistatic interactions that are physically close in the genome and likely inherited together.

The most widely studied population genetic models, those for eukaryotes, have not been directly applicable to bacteria (or Archaea; *13*), and consequently we know little about how epistasis affects adaptive processes and the evolution of multi-locus phenotypes such as antibiotic resistance, antigenic profile, or metabolic output, all of which likely involve epistatic interactions (*14*–*16*). Answering these questions, which are relevant for the entire bacterial and Archaeal kingdoms of life, requires multi-locus models with epistasis that account for the unique features of bacterial recombination that involves the transfer of smaller DNA segments, together with accurate estimates of genome-wide recombination parameters.

We develop a multi-locus model of bacterial evolution to study how selection may act on standing genetic variation when both additive and epistatic effects contribute to fitness differences between individuals. In this model, we vary the magnitude and genetic basis of fitness differences but use biologically realistic levels of bacterial recombination by inferring these parameters from genomic data of five pathogens (*Staphylococcus aureus, Campylobacter jejuni, Streptococcus pneumoniae, Neisseria gonorrhoeae*, and *Helicobacter pylori*), using Approximate Bayesian Computation (ABC) and machine learning. The bacteria we chose exhibit strikingly different degrees of genome-wide LD and include some of the most highly recombining bacteria known. In contrast with eukaryotes, we find that in bacteria selection may act on the cumulative effects of very weak epistatic interactions (N|s| ≈ 1-10) regardless of their physical proximity on a circular chromosome and does not require specific metapopulation structures that have historically been invoked by Wright and others to limit the impact of recombination (*5*,*6*), even for highly recombining pathogens that have been previously labeled as freely recombining (*12*). Thus, while recombination is sufficiently strong in many bacteria to destroy phylogenetic signal in gene trees (*17*) and to prevent periodic selective sweeps from purging genome-wide variation (*18*,*19*), it is not capable of hindering selection on epistatic interactions between polymorphisms.

## Results

### Recombination parameter estimates

In order to simulate bacterial evolution with selection and biologically realistic recombination rates, we first inferred recombination parameters using five genomic datasets from *S. aureus, C. jejuni, S. pneumoniae, N. gonorrhoeae*, and *H. pylori*. We inferred both the rate of DNA transfer and the mean tract lengths involved using a new approach that we developed (below), as opposed to using previously published estimates, because other popular recombination-detection programs have known biases towards detecting larger recombination events between more diverged sequences (*20*). While these programs may miss short recombination events or transfers between less diverged sequences, these events affect pairwise compatibility (PC) and linkage disequilibrium (LD) such that use of these summary statistics facilitates parameter inference in species that have less diversity (e.g. *N. gonorrhoeae*) or exchange short DNA tracts. Consequently, methods that use correlations between mutations to quantify recombination have been gaining popularity (*21*,*22*). Both the rate and mean lengths of DNA transfers have critical implications for how selection acts on epistatic interactions.

For genomic datasets containing isolates collected across many years or large geographic areas, we selected a restricted subsample (Table S1) to avoid population structure that could confound estimates of recombination. Analysis of samples taken at very different time points may artificially elongate terminal branches of the genealogy, leading to underestimates of LD (and related statistics) and overestimates of recombination (*23*). On average, each dataset had over 1,000 core genes containing almost 1Mb of DNA (Table S2). We used fourfold degenerate sites in each sample to calculate Tajima’s D, which was typically near zero (Table S2), suggesting these samples came from populations that have not experienced nonequilibrium demography and are not strongly structured, both of which may affect estimates of recombination parameters. More information on data processing can be found in the Materials and Methods.

Using Approximate Bayesian Computation (ABC) coupled with Bayesian Optimization (*24*), we fit customized coalescent models with gene conversion to summaries of genomic data in order to infer three parameters: the population mutation rate *θ*=2*Nμ*, the population recombination rate *ρ*=2*Nr*, and the mean of DNA tract lengths transferred between donor and recipient (~Geometric(*q*), where 1/*q* is the mean tract length). To summarize the statistical associations between single-nucleotide polymorphisms (SNPs), we developed an approach that gave accurate estimates of *ρ* and *q* (Materials and Methods). Briefly, we used PC to quantify the amount of recombination that has taken place between two SNPs, which are compatible with a single phylogeny if fewer than four haplotypes are observed (*25*); either recurrent mutation or, more likely, recombination gives rise to four haplotypes (Figure S1). PC is equivalent to the four-gamete test (*26*) and quantifies historical recombination similar to measures of LD, such as D’ or *r*^2^ (*27*). We quantified PC within *k* genomic windows and compared these observed estimates to *k* simulated windows via a Kolmogorov-Smirnov statistic, which captures higher moments of the PC distribution (see Figure S2 for a diagram of our analysis). To infer *θ* we calculated the minimum number of mutations per site (*28*), a sufficient statistic for this parameter (*29*). With these summary statistics, input recombination parameters were accurately recovered from simulated datasets (Figure S3, S4).

Recombination estimates for the 5 bacteria studied are summarized in Table 1, and simulations with these parameters largely fit observed data (Figure 1). While our parameter estimates for some species (*S. pneumoniae* and *C. jejuni*) are generally consistent with previous work, we observe some differences for other species that could be relevant to selection on epistatic interactions. For instance, parameter estimates for *H. pylori* revealed an extremely high value of *ρ*=472 per kb but short tract lengths around ~50 bp that are approximately an order of magnitude smaller than previous estimates of ~400 bp from genomic data (*30*,*31*), yet in agreement with short lengths reported in carefully designed *in vitro* experiments that used diverged donor and recipient strains (*32*). We also find that *S. aureus* frequently transfers (*ρ*=11.5 per kb) tracts around ~70 bp in length, which are also an order of magnitude smaller than previous estimates of ~650 bp (*33*). Nonetheless, *S. aureus* still exhibits high genome-wide linkage, since these small transfers affect few SNPs. For both species, these short tract lengths agree with PC decaying within 100bp (Figure 1), as the PC vs. distance distribution is expected to asymptote near the mean tract length because SNPs separated by larger distances are equally likely to be unlinked. Such short recombination events have important consequences for evolution (below).

**Figure 1.**
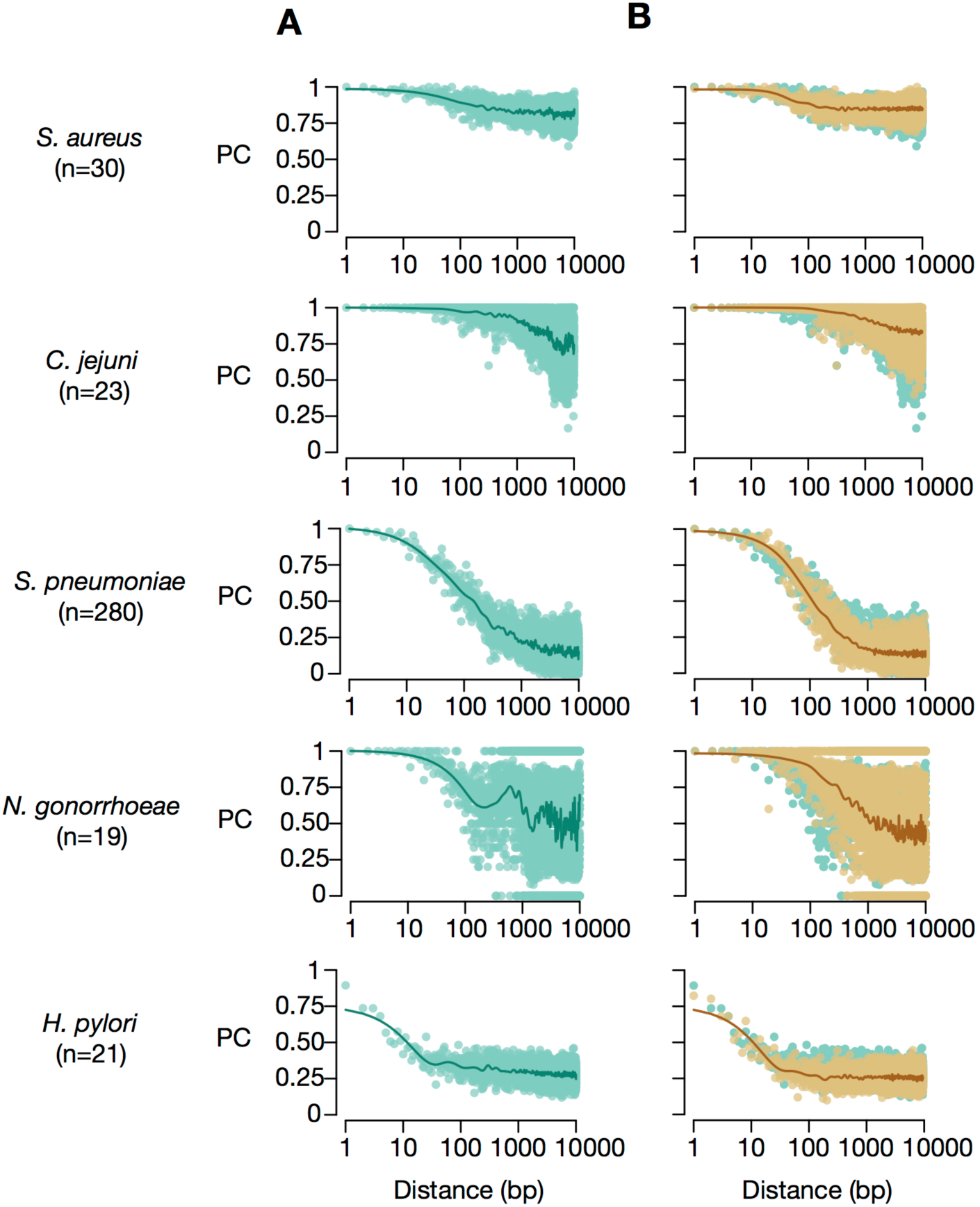
Observed pairwise compatibility vs. distance. **(A)** Patterns of PC (green) vary among the species included in this study. Since PC measures the compatibility of two SNPs with a single phylogeny, these data indicate that SNPs more than ~1kb apart have distinct phylogenetic histories from recombination, with the exception of *S. aureus* which exhibits linkage. **(B)** Simulated patterns of PC vs. distance (brown) using parameter estimates from Table 1 fit observed data well. We note that the sensitivity of PC to sample size (n) makes these patterns not directly comparable across species and that the product of effective population size and recombination (or *ρ*=*2Nr*) affects PC.

**Table 1.** ABC Parameter estimates

| Species | Parameter | MDE | Mean<br>(Lower CI/Upper CI) |
| --- | --- | --- | --- |
| ***S. aureus***<br>(n=30) | $\theta$ | 31.9 | 32.0 (24.6/41.6) |
| | $\rho$ | 11.5 | 12.2 (6.9/22.2) |
| | $1/q$ | 68 | 63 (30/120) |
| ***C. jejuni***<br>(n=23) | $\theta$ | 10.6 | 10.6 (7.6/14.4) |
| | $\rho$ | 0.6 | 0.6 (0.3/1.0) |
| | $1/q$ | 1429 | 1429 (667/3333) |
| ***S. pneumoniae***<br>(n=280) | $\theta$ | 26.9 | 26.7 (19.8/35.5) |
| | $\rho$ | 11.5 | 11.3 (9.2/13.9) |
| | $1/q$ | 588 | 625 (455/833) |
| ***N. gonorrhoeae***<br>(n=19) | $\theta$ | 4.3 | 4.9 (1.4/19.5) |
| | $\rho$ | 8.6 | 8.2 (4.5/14.5) |
| | $1/q$ | 2500 | 3333 (1250/10000) |
| ***H. pylori***<br>(n=21) | $\theta$ | 133.7 | 129.0 (106.1/160.5) |
| | $\rho$ | 471.7 | 484.7 (378.8/652.7) |
| | $1/q$ | 49 | 49 (27/96) |

Note: MDE represents the minimum discrepancy estimate from the GPR, or the parameter value set with the smallest discrepancy between simulated and observed data. Estimates of $\theta$ and $\rho$ are in units per kb, while $1/q$ is in bp.

To our knowledge, this analysis is the first to infer both a recombination rate (*ρ*=8.6 per kb) and mean tract length in *N. gonorrhoeae*, which appears to transfer long segments (~2.5 kb) since PC decays slowly with distance (Figure 1). The PC data is notably noisy (in particular the change that occurs among SNP pairs separated by ~500-700bp), which may be due to rearrangements that have occurred since the divergence of our sample and the reference sequence used to estimate inter-SNP distances. Results are similar when we use a different reference sequence (Figure S6A), suggesting the rearrangement may be a derived feature of our sample. While we do not know the exact effect this noise has on our parameter estimates, it may have led to a slight overestimate of recombination rates, as mean PC from simulations parameterized with MDE values listed in Table 1 is lower (~10%) than mean PC between randomly sampled SNP pairs, irrespective of distance (Figure S6B).

We thus have within our dataset parameter estimates from a diverse set of bacteria that represent the many ways bacteria may transfer DNA, including very high or low rates of transfer with short or long tract lengths. A full description of the parameter inference results can be found in the Supplementary Text.

### Multi-locus simulations of evolution

We used simulations of bacterial evolution to study how epistasis may affect the short-term rate of adaptation, measuring evolutionary time via changes in diversity arising through selection and genetic drift. We model *L* loci arbitrarily located on a circular chromosome (following Fisher’s infinitesimal model). The recombination parameter estimates obtained above are coupled with previously published mutation rates (*μ*) to derive the physical recombination rates (*r*) with *r*=*ρμ*/*θ* (Table S3). We vary the total variance in fitness (*σ*^2^), which captures the magnitude of fitness differences between individuals, and the genetic basis of *σ*^2^, which is comprised of an additive component (*V_A_*) from individual locus effects and an epistatic component (*V_I_*) from pairwise interaction effects, with *σ*^2^=*V_A_*+*V_I_*. Our model uses sign epistasis, with effect sizes drawn from a normal distribution with mean zero (Materials and Methods). Populations were initiated with standing genetic variation (all *L* loci polymorphic) and evolved until diversity decayed by 20%, as measured by the average number of pairwise differences. At this point, we recorded patterns of LD and population fitness relative to an asexual control with the same epistatic effect sizes to directly compare our results to those expected under clonal evolution. For contrast, we also modeled a linear eukaryotic chromosome with a relatively low crossover recombination rate equivalent to facultatively sexual yeast (Table S4).

With 10 loci randomly distributed along a ~2Mb chromosome, simulations with eukaryotic recombination rates exhibited similar short-term responses to selection compared to an asexual control, when population variance in fitness was primarily due to additive effects (Figure 2A). However, if the same selective differences were due to epistasis, the response to selection diminished dramatically, showing that even low levels of crossover recombination may antagonize adaptation. Bacterial simulations, on the other hand, had faster rates of adaptation. Simulations with parameters from *H. pylori* behaved similar to eukaryotes for highly epistatic traits (*V_I_*/*σ*^2^ = 1) composed of very weak interactions (*N|s|* ≈ 0.1-0.4, front right) but these dynamics quickly changed once interactions became moderately strong (*N|s|* ≈ 4-10) (Figure 2B). Theory from single-locus population genetics has shown selection efficiently acts on mutations only when *N*|*s*| >>1 (*34*). A hallmark of this clonal selection regime is increased LD measured as D’ between selected loci (Figure 2E; *35*,*36*). Selection responses were also higher for *H. pylori* simulations when epistasis only contributed an intermediate amount to the variance in fitness (*V_I_*/*σ*^2^ ≥ 0.5) and selection was sufficiently strong (back rows in Figure 2B). *S. pneumoniae*‘s response to selection was relatively insensitive to the degree of epistasis *V_I_*/*σ*^2^ for much of the parameter space explored here.

**Figure 2.**
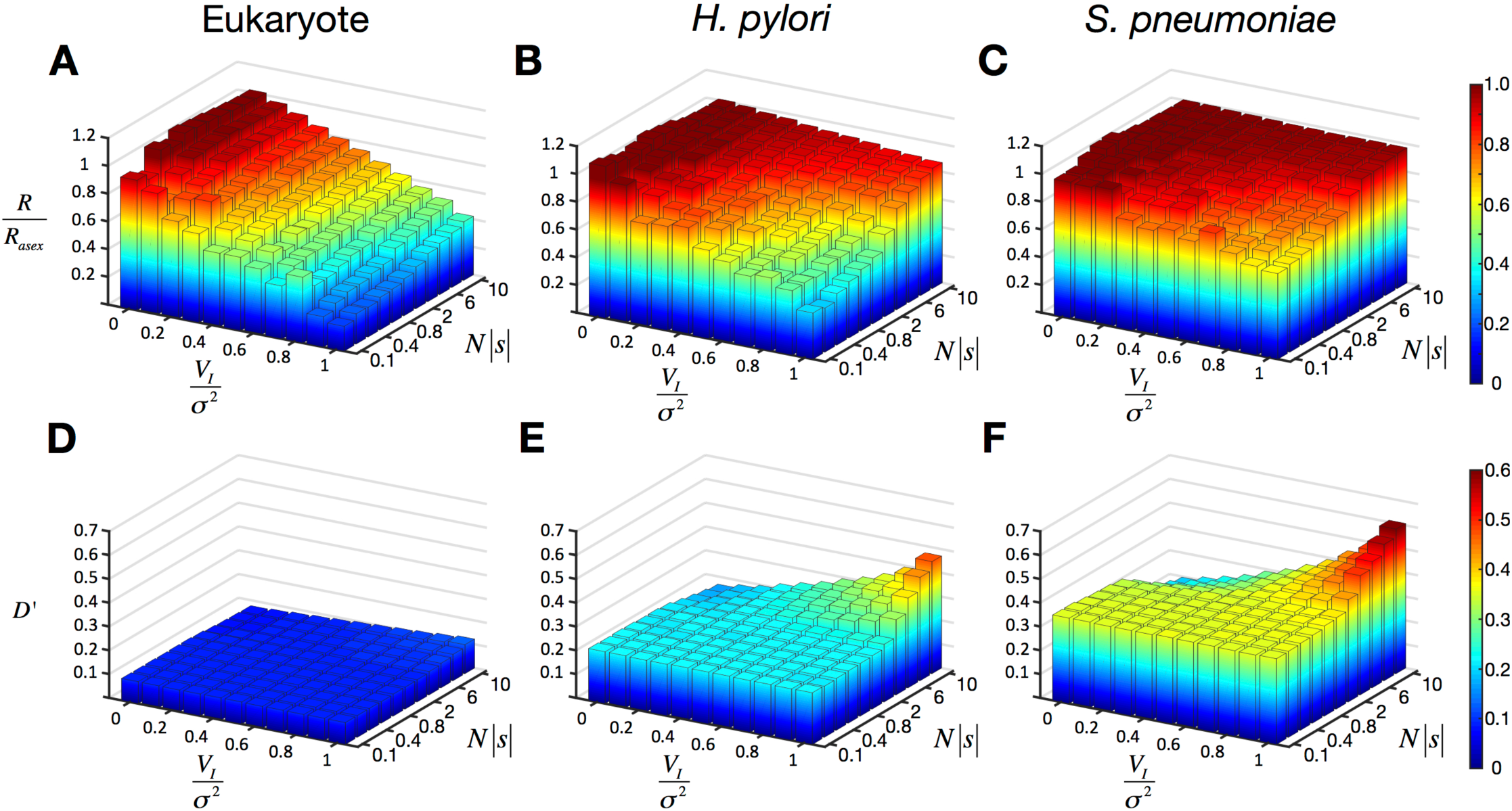
Bacterial recombination interferes with the response to selection, especially for highly epistatic traits under weak selection, but less than eukaryotic recombination. Response to selection, relative to an asexual control (*R/R_asex_*) for a eukaryote (**A**), *H. pylori* (**B**), and *S. pneumoniae* (**C**). The relative epistatic variance (*V_I_*/*σ*^2^) varies along the x-axis, and the variance in fitness (*σ*^2^) increases along the y-axis but is labeled according to the mean value of *N|s|* for each pairwise epistatic interaction when *V_I_*/*σ*^2^ = 1. (**D-F**) Linkage disequilibrium, as measured by D’, increases when selection outcompetes recombination and spreads particular genotypes through the population. The result for each parameter set is an average of 1000 simulations using L=10 loci under selection, randomly distributed throughout a genome.

The short-term selection responses for simulations with recombination parameters estimated for other bacterial species were similar when the same 10-locus trait was fully or partially epistatic (Figure 3A, B respectively), and all bacterial rates of adaptation were consistently higher than the eukaryotic model (colored lines vs. black lines). However, if these 10 loci were instead randomly distributed within 5kb, (such as might be the case for an operon) selection efficiently acted on epistatic effects in the eukaryote (Figure 3C,D; black line), as expected, since recombination breakpoints occur less frequently within smaller intervals. In bacteria, this distance-dependence in the effectiveness of selection depended on the mean tract length of DNA transferred between donor and recipient. For example, *H. pylori* transfers ~50bp fragments that are similarly unlikely to unlink SNPs separated by 1kb or 1Mb. Alternatively, *N. gonorrhoeae* has longer recombination tract lengths that more likely co-transfer SNPs within 2.5kb, the mean tract length. While these parameters allowed selection to more efficiently act on physically close loci, the effect was weaker than that observed in the eukaryotic model (Figure 3). Irrespective of tract lengths, these results show the overall sensitivity of bacterial evolution to epistasis between polymorphisms depended on the fraction of bases transferred each generation via homologous recombination (Table 2), which we refer to as *F_rec_*.

**Figure 3.**
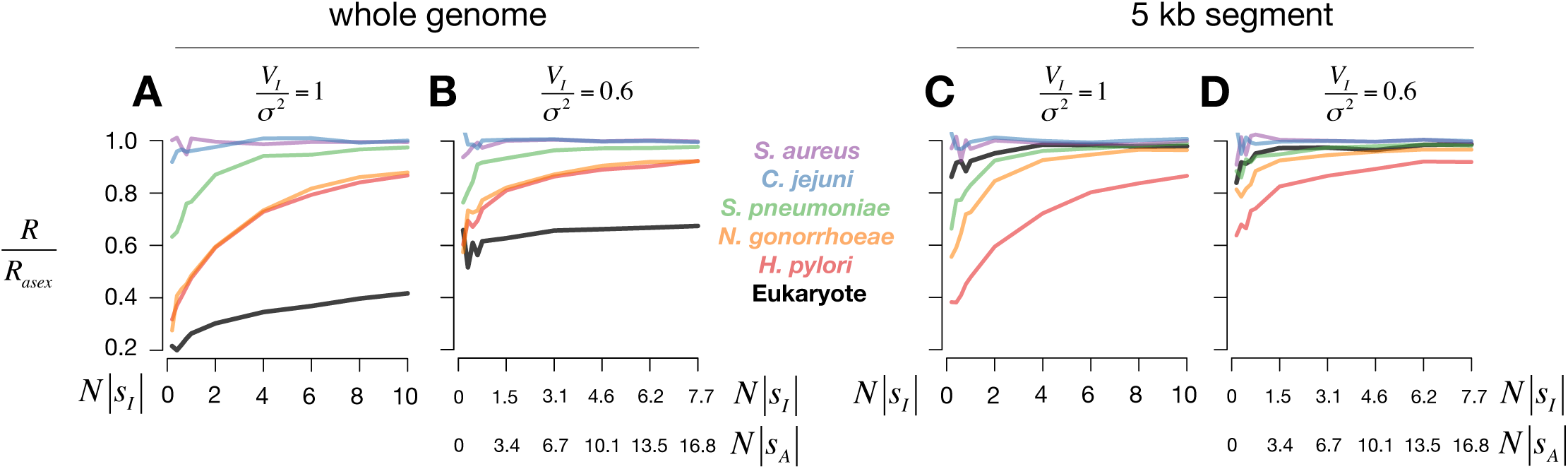
Distance-dependence of selection on epistatic interactions. Short-term relative selection responses are shown for a trait controlled by *L*=10 loci randomly distributed throughout a genome **(A,B)** or 5kb segment **(C,D)** when the variance in fitness is purely or partially epistatic (*V_I_*/*σ*^2^ = 1 or 0.6, respectively). The epistatic effect per locus pair (*N|s_I_*|) and additive effect per locus (*N|s_A_*|) are shown along the x-axis. Colored lines correspond to different bacterial species. As in Figure 2, **A** and **B** show selection acts more efficiently on epistatic traits in bacteria than in the eukaryote (black line). While selection acts significantly more on physically close epistatic interactions for the eukaryote **(B,D),** this effect is not as strong for bacteria, particularly for species that transfer short tract lengths (*H. pylori*). Increasing the amount of additivity **(B,D)** makes all species have greater selection responses since fitness depends less on particular allele combinations and is not altered as much by recombination events.

**Table 2.** Fraction of bases exchanged per generation via homologous recombination (*F_rec_*)

| Species | $\frac{\rho}{q(\text{Genome Size})}$ |
| --- | --- |
| *S. aureus* | $0.28 \times 10^{-06}$ |
| *C. jejuni* | $0.52 \times 10^{-06}$ |
| *S. pneumoniae* | $3.24 \times 10^{-06}$ |
| *N. gonorrhoeae* | $9.98 \times 10^{-06}$ |
| *H. pylori* | $13.9 \times 10^{-06}$ |

\*recombination parameter values from MDEs in Table 1, genome sizes from Table S5

Simple traits may be controlled by fewer loci (e.g. compensatory mutations that ameliorate the cost of antibiotic resistance), so we also modeled a trait with 3 loci using the same additive and epistatic selection coefficients as above (Figure 4). With fewer loci, *σ*^2^ decreases (fewer additive effects and interactions per individual), increasing the relative amount of recombination to selection (*r*/*σ*^2^). Consequently, models of the most recombining bacteria (*H. pylori* and *N. gonorrhoeae*) exhibited diminished selection responses for highly epistatic traits with weak effects, although responses increased within the range *N|s|* ≈ 2-10, while the eukaryote responses changed little. Efficient selection on simple epistatic traits in these species may thus frequently involve larger interaction effect sizes (*N|s|* > 10), which is likely true for antibiotic resistance and compensatory mutations that often have very large effects (*37*). In contrast, simulations mimicking highly recombining *S. pneumoniae* still behaved in a largely clonal fashion (*R/R_asex_* ≥ 0.6) for even weak effects. *S. aureus* and *C. jejuni* models did not appear to recombine frequently enough to compete with even weak selection and had similar responses to asexuals for any number of loci or effect sizes examined here (Figure 4).

**Figure 4.**
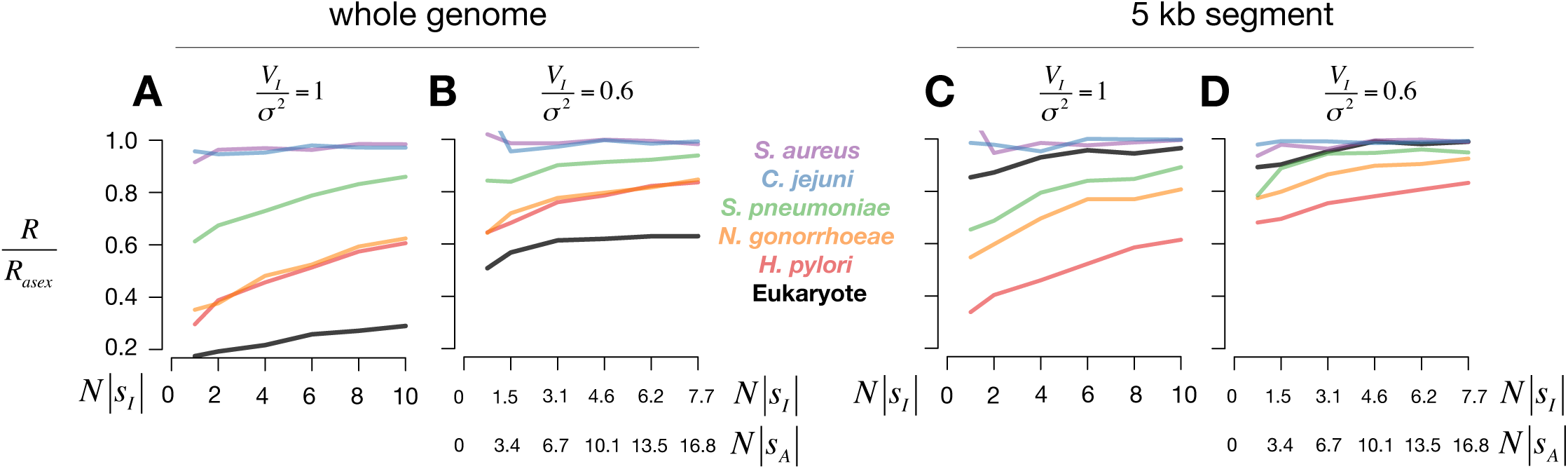
Selection responses for a less complex trait. As in Figure 3, short-term relative selection responses are shown for a trait controlled by *L*=3 loci randomly distributed throughout a genome **(A,B)** or 5 kb segment **(C,D)** when the variance in fitness is purely or partially epistatic (*V_I_*/*σ*^2^=1 or 0.6, respectively). While the responses here are similar to the previous case of 10 loci (see Figure 3 for more details), the magnitude is diminished because for a given set of additive and epistatic effect sizes (x-axis), *σ*^2^ decreases with the number of loci. Note that data are plotted only for *N|S_I_|* > 1 as simulations with smaller *σ*^2^ were highly stochastic.

### Distributions of pairwise LD between synonymous SNPs

Our finding that even high rates of bacterial recombination permit epistatic alleles to contribute to adaptation raises the question of whether epistasis has shaped patterns of PC, which we used to quantify recombination under a neutral model. While we attempted to mitigate the affect of selection on parameter estimates by using synonymous SNPs, these polymorphisms could theoretically exhibit epistatic interactions or be partially linked to epistatic nonsynonymous SNP pairs, both of which may skew distributions of LD (and thus PC; *38*) and lead to underestimates of population recombination rates if we assume these polymorphisms are neutral. However, whether or not epistasis affects genome-wide patterns of LD depends on the frequency and strength of epistatic interactions and whether selection is ongoing in the population (i.e. loci under selection have not fixed).

To assess whether epistasis has recently shaped the statistical associations between synonymous SNPs, we polarized mutations using an outgroup species (Table S6) and quantified LD using *D* = *p_ij_-p_i_p_j_*, where *pi* and *p_j_* are the derived allele frequencies at site *i* and *j*, and *p_ij_* is the observed frequency in which both derived alleles occur on the same haplotype. The expected value of *D* is zero under neutrality, but epistasis or population structure can skew the distribution away from this expectation, even if epistatic fitness effects are equally likely to have positive or negative values (*38*). For most species examined here, observed distributions of *D* between synonymous SNP pairs largely resembled neutral distributions simulated with the MDE parameters inferred in Table 1 (Table S7). Observed distributions of *D* had slightly different means (all near zero) but similar variances compared to neutral simulations (Table S7), suggesting epistasis has not strongly shaped genomewide LD between synonymous SNPs. However, we cannot completely rule out selection having small effects on parameter estimation. Incorporating selection into simulation-based models for parameter inference requires knowledge of the distribution of epistatic effects, which we are currently exploring.

## Discussion

Simple evolutionary models from population genetics shape our expectations and interpretations of nature and guide the development of future research (*3*,*5*,*9*). Interest in sexual eukaryotes has inspired the simplifying convention of gene-centered models that promote the importance of additive gene effects and ignore epistasis; multilocus genotypes in eukaryotes rarely persist longer than a single generation such that only anomalously strong interactions require consideration (*39*). This gene-centric view pervades discussions of evolution and adaptation (3–7), including those on bacteria, many of which recombine extensively and are commonly reduced to “core” and “accessory” genes, with niches providing selective advantages to specific genes (*40*,*41*). While this model is certainly inappropriate for largely clonal bacteria, we show, in the presence of epistasis, it also may not apply to highly recombining bacteria thought to be “effectively sexual” (*11*) from very low correlations between SNPs (*r*^2^, Figure S7), even if the epistasis is weak (*N|s|* ≈1-10) and loci are distantly spaced. Selection in many bacteria may thus act on epistatic interactions that would be virtually invisible to selection in eukaryotes, allowing bacteria to harness these effects to rapidly respond to novel pressures. Selection may act on even weaker epistatic effects for fitness traits controlled by more than 10 loci (the maximum explored here), since increasing the number of loci with fitness effects increases *σ*^2^ and thus decreases *r*/*σ*^2^ (*36*). The stark difference in the ability of epistatic alleles to contribute to adaptation in bacteria and eukaryotes is driven by differences in genome architecture. Crossover recombination in eukaryotes exchanges large genomic segments, breaking numerous allele pairings even under low rates of exchange. Homologous recombination, alternatively, involves the transfer of shorter DNA segments that can substantially reduce LD between nearly neutral polymorphisms over time (Figure S7) but less so over the shorter evolutionary timescales on which selection acts.

These findings generate experimentally testable predictions relevant to exploring the distribution of epistatic effects between natural polymorphisms: 1.) effect sizes (in terms of *Ns*) will generally be stronger for species that recombine extensively, for instance compensatory mutations for antibiotic resistance may be stronger in species such as *N. gonorrhoeae* and *H. pylori*, or even less likely to exist; 2.) the distribution of observed epistatic effect sizes may differ between polygenic fitness traits controlled by fewer or more loci since, for a given amount of recombination, selection may act on weaker epistatic effects if there are more loci contributing to fitness differences; 3.) short range interactions between physically closer loci may be broken more frequently in highly recombining species that exchange short DNA tracts (e.g. *H. pylori*), which may alter the distribution of observed epistatic effects between loci within operons. Our results also have important implications for the genetic basis of adaptation and the maintenance of population structure, since epistatic alleles may be maintained even in the presence of high levels of homologous recombination.

We have quantifed bacterial recombination parameters with a new method sensitive enough to detect small tract lengths and events between closely related strains. We summarize these recombination parameters as *F_rec_* (Table 2), which largely dictates the ability of selection to act on epistasis between distant polymorphisms (Figure 3A,B). For physically close pairwise interactions within 5kb, bacteria may decouple allele combinations faster than eukaryotes, particularly for species with higher recombination rates and shorter tract lengths (Figure 3C,D). Consequently, the evolutionary benefit to physically clustering loci that epistatically interact may be weaker in bacteria (close and distant loci are decoupled with similar frequencies) than eukaryotes, in which epistasis is thought to play a role in gene order (*42*).

We use sign epistasis in our simulations (Materials and Methods), which was also used in historical arguments to challenge Fisher on the potential importance of gene interactions (*5*,*10*). While the specific type of epistasis will have important consequences for long-term evolution, we have focused on the short term to define the fundamental capacity of selection to act on allele combinations in bacteria. Ultimately, the importance of epistasis will also depend on the distribution of epistatic fitness effects, which may vary between traits, and the number of loci that contribute to multi-locus fitness traits. Recent work in *S. pneumoniae* revealed evidence of co-evolution across the genome, with numerous loci exhibiting elevated LD (*43*), some of which may be driven by selection on epistatic interactions as our results here show it is very plausible for selection to act even on weak epistasis in this species (Figures 2-3).

While our dataset consists of opportunistic and obligate pathogens, our results likely extend to other microbes. For instance, a genomic study of *Vibrio cyclitrophicus* showed recombination is sufficiently strong to allow gene-specific selective sweeps, as opposed to the periodic selection model in which sweeps have genomewide effects (*19*). Like in *H. pylori*, LD decays rapidly within 50 bp in *V. cyclitrophicus* but asymptotes at a value well above zero, suggesting residual linkage genome-wide. Thus, while gene flow and recombination may homogenize ecotypes outside of genomic regions involved in local adaptation, recombination may not be strong enough to antagonize selection on epistatic interactions. However, exact estimates of recombination from LD data require knowledge of population size (via knowledge of the mutation rate). We also applied our method to a genomic dataset of thermophilic archaea *Sulfolobus islandicus* (*44*) but struggled to accurately infer the mean tract length due to low diversity and small sample size (11 isolates). Nevertheless, our best parameter estimates strongly suggest a low recombination rate that likely permits selection on weak epistatic interactions (Figure S8). Thus, epistasis between polymorphisms scattered across the genome may play a critical role in adaptive processes across a majority of the tree of life, and unlike eukaryotes, these interactions do not need to be strong or physically close, and do not require specific metapopulation dynamics to permit efficient selection.

## Materials and Methods

### Genomic data

We analyzed previously published genomic datasets (Table S1). Using an amino acid file from a reference genome (Table S2), we annotated *de novo* assemblies from each species with PROKKA (*45*) and identified core and accessory genes with ROARY (*46*). Only core genes ‐‐ present in all samples in a species – that were also present in the reference genome were used for downstream analyses. All position information between genes and polymorphic sites is derived from their relative positions in the reference genome used to annotate *de novo* assemblies, not from a reference-based DNA alignment (Figure S2). For analyses that required polarized mutations, we used Mauve (*47*) to align these reference genomes to an outgroup species (Table S6) to infer the derived and ancestral state of each polymorphism.

### Sub-sample selection

For datasets with a wide geographic or temporal distribution, we partitioned samples by geography and collection date into smaller subsamples to minimize the effects of sampling and structure on population genetic parameter estimates. Subsamples consisted of isolates from a similar geographic region (to avoid genetic isolation by distance) and also had similar collection dates, since serial samples can skew genealogies to have longer terminal branches, potentially leading to substantial overestimates of linkage disequilibrium or related statistics (*48*). Subsamples that had the least evidence of substructure (near-zero estimates of Tajima’s D) were chosen for analysis (Table S2).

### Estimation of population genetic parameters

#### Summary statistics

We used five summary statistics to fit coalescent models to observed genomic data: the minimum number of mutations per site (*28*) to estimate the population mutation rate *θ*=2*Nμ*, and four recombination-related statistics to estimate both the population recombination rate *ρ*=*2Nr* and the mean of the geometrically-distributed DNA tract lengths transferred between donor and recipient (1/*q*, where *q* is the geometric distribution parameter). We used pairwise compatibility (PC) to quantify the amount of recombination that has taken place between two SNPs, which are compatible with an infinite-sites model of no recombination if less than four haplotypes are observed; either recurrent mutation or, more likely, recombination gives rise to four observed haplotypes (Figure S1). PC generally decreases as a function of the genomic distance between SNPs in recombining bacteria, and the shape of this decay contains information about both *ρ* and q. Consequently, for four different inter-SNP distance categories we calculated mean PC to capture both short- and long-range recombination dynamics (similar to (*21*)). We calculated mean PC only for synonymous, intermediate-frequency SNPs (10-90% frequency in sample), and the minimum number of mutations only for fourfold degenerate sites.

Since we are interested in inferring parameters genome-wide, we compared observed summaries from *k* genomic windows with *k* simulated datasets, since our coalescent model could only simulate segments of maximal length roughly equal to 150 kb (below). To find the right set of summary statistics for inference of *ρ* and *q*, we simulated a *k*-window dataset with known parameter values and compared summaries calculated from this “true” dataset with those calculated from k-window datasets simulated across a grid of *θ*, *ρ*, and *q* values. Specifically, for each simulated dataset in the grid, we calculated a discrepancy between *k* simulated and “true” summary statistic values using a Kolmogorov-Smirnov statistic, one for each of 5 summaries (above). We summed these 5 Kolmogorov-Smirnov statistics for each point in the grid (Figure S3). When we measured PC at two short-range inter-SNP distances (with respect to the decay of PC vs. distance) and two long-range inter-SNP distances, datasets simulated with parameter values close to “true” values had the smallest discrepancy. Thus, we chose different inter-SNP distances and window lengths for each species (Table S5) based on the decay of PC vs. distance and on the computational resources needed to simulate data within the parameter space, as high *ρ* required more memory or time, depending on the simulator used. We found that only comparing the mean values of PC summaries between simulated and observed data, as opposed to the full distribution of *k* values, resulted in a reduced ability to distinguish between datasets simulated with high *ρ*, large *q* (small tract lengths) and low *ρ*, small *q* (large tract lengths).

#### Coalescent simulations

Using CoaSim (*49*), we constructed a novel, finite-sites coalescent model to simulate genomic DNA, which was required to accurately model sequence alignments from highly diverse pathogens like *H. pylori* that frequently have multiple bases per site. A finite-sites model not only enables more accurate inference of *θ*=2*Nμ* but also more precise estimates of recombination parameters since back mutation mimics recombination by affecting PC, particularly for species with high transition:transversion ratios (Ti:Tv). Our coalescent model thus accounted for species-specific base compositions and Ti:Tv, which we estimated from a reference genome or fourfold degenerate sites, respectively (Table S5). For less diverse species, we used ms (*50*) to simulate longer DNA sequences, which yielded better estimates of recombination parameters as there were more SNP pairs with particular inter-SNP distances.

#### Parameter Estimation

We constructed a novel approach to estimate population genetic parameters in bacteria, based on the statistical framework of (*24*). Our method uses Bayesian Optimization with Gaussian Process regression (GPR) to model the discrepancy between simulated and observed datasets. We chose this machine-learning approach to prudently explore parameter space due to the computational requirements of the finite-sites coalescent simulator. For each set of parameters (*θ*, *ρ, q*) chosen by the method, we simulated *k* genomic windows, calculated 5 summary statistics (above) for each window, and used a Kolmogorov-Smirnov statistic to calculate 5 discrepancies between each set of *k* simulated and observed summaries. The final discrepancy was a sum of these 5 Kolmogorov-Smirnov statistics. We ran the inference algorithm for 320 iterations, simulating *k* windows each time (Figure S5). The results stabilized after 200 iterations when the GPR no longer changed with additional acquisitions (visual inspection). The GPR model of the discrepancy was then used in an approximate Bayesian computation (ABC) framework to approximate the intractable likelihood function that enabled us to compute the posterior distributions by standard sequential Monte Carlo sampling (*24*). Specifically, the likelihood function was approximated by the probability to draw discrepancy values from the GPR model that are less than a small threshold. Following common practice, the threshold was set to the 1% quantile of the discrepancy values of the simulated data. When given “observed” data that we generated with known parameter values, our approach accurately estimated the input parameter values (Figure S4).

Comparing *k* simulated DNA segments with *k* observed genomic windows implicitly assumes independence among windows, since each coalescent simulation is independent. To test whether this assumption affects parameter estimation, we simulated a large 200kb DNA segment with a modest recombination rate and broke this segment up into 10 20kb windows. We then simulated a grid of *θ*, *ρ*, and *q* values around the “true” values used to simulate the large segment and calculated the discrepancy between simulated and “true” summaries, as above. Like before, datasets simulated with parameter values close to “true” values had the smallest discrepancy (Figure S3), showing that this independence assumption should not greatly affect our parameter estimates and conclusions, at least within the range of recombination rates studied here.

### Epistasis simulations

#### Model

We created a forward-time simulation framework in C++ similar to that of Neher and Shraiman (*36*), which uses Fisher’s infinitesimal model to simulate a population of chromosomes with *L* loci represented as *l_i_* = ±1, *i* =1, … *L*. Each locus has an additive effect 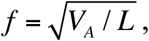 where *V_A_* is the additive fitness variance, and each locus pair has an epistatic interaction effect *f_ij_* ~ *Normal*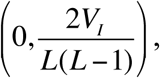 where *V_I_* is the epistatic fitness variance. Distributions of epistasis with zero mean have been observed in bacteria and other microbes (*2*). The fitness of each individual is calculated as:

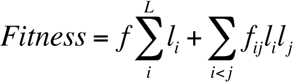
such that each locus or locus pair contributes a small, similar amount to the additive or epistatic component of fitness, respectively. However, we modified Neher and Shraiman’s framework in two important ways: (*1*) we modeled both homologous recombination in a circular chromosome and eukaryotic recombination on a linear chromosome and (*2*) we randomly distribute the *L* loci in a circular/linear segment of arbitrary size, allowing us to accommodate for different bacterial genome sizes but also simulate loci within smaller sub segments (e.g. 5kb). While we simulate recombination genome-wide, only those events that overlap with any positions of the *L* loci affect the genotype, unless donor and recipient have the same alleles at those loci. Source code for this simulator is freely available (github.com/brian-arnold/BacteriaEpistasisSimulator).

#### Analysis of simulations

We ran each simulation until genetic diversity (as measured by the average number of pairwise differences π (*51*)), decayed a specified amount in order to analyze simulations of different population sizes on the same evolutionary timescale. At this stopping point, we calculated the standardized response to selection

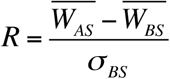
or the difference between the mean population fitness before 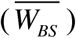 and after 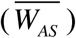 selection.

We also ran an asexual “control” for each simulation, holding all parameter values constant (even pairwise-epistatic selection coefficients, which are randomly drawn for each replicate of bacterial evolution) but setting the recombination rate to zero, and calculated the same response to selection (R_asex_). We used these two quantities to calculate the relative speed of adaptation *R/R_asex_* and study how clonal these pathogens behave in the presence of varying amounts of epistasis. For short-term evolution (e.g. π decays to 80% starting value), *R/R_asex_* <= 1 if much of the variance in fitness is epistatic, since recombination will primarily break apart beneficial genotypes. For long-term evolution or if fitness effects are primarily additive, *R/R_asex_* may be >= 1 from greater exploration of the fitness landscape.

#### Rescaling simulations

We inferred both the population mutation rate *θ*=2*Nμ* and population recombination rate *ρ*=*2Nr* from genomic data, but in order to obtain estimates of the recombination rate (*r*) that we used in simulations with epistasis, we used previously published mutation rates (*μ*) to calculate *r*=**ρ*μ*/*θ*. However, these mutation rates come from serial samples and are scaled in units of months or years, not the standard generation timescale normally used in Wright-Fisher models. Consequently, by using these scaled estimates of *N* and *r*, we effectively simulate a rescaled population with *Ν*/*λ*, *λ*r, and *λs*, where *λ*>*1* represents the number of bacterial generations per time step (i.e. 1 month). While rescaling has been previously explored for simple scenarios of selection (*52*), we confirm that rescaling preserves the population genetic dynamics of more complex selection with epistasis and recombination as long as the ratio of *r/s* is conserved. For instance, simulating a population size of *N* = 10,000 or *N/λ* = 1000 (λ=10) gives remarkably similar results on different evolutionary timescales (different by a factor of λ), as long as *r* and s are changed accordingly such that *Nr, Ns*, and *r/s* are constant (Figure S9).

## Acknowledgements

All data used in this study came from previously published papers shown in Table S1. BJA was supported by a postdoctoral fellowship from the National Institute of Health (1 F32 GM120839-01). The authors declare no competing financial interests. The computations in this paper were run on the Odyssey cluster supported by the FAS Division of Science, Research Computing Group at Harvard University.

## Author Contributions

BJA, WPH, and ML conceived and headed the project. BJA, MUG and JC designed the statistical framework used to infer recombination parameters. BJA performed all simulations and bioinformatic analyses. BJA, WPH, and ML wrote the manuscript with input from all coauthors. SKS provided data for *Campylobacter jejuni*.

